# Factors influencing gene family size variation among related species in a plant family

**DOI:** 10.1101/270009

**Authors:** Peipei Wang, Bethany M. Moore, Nicholas L. Panchy, Fanrui Meng, Melissa D. Lehti-Shiu, Shin-Han Shiu

## Abstract

Gene duplication and loss contribute to gene content differences as well as phenotypic divergence across species. However, the extent to which gene content varies among closely related plant species and the factors responsible for such variation remain unclear. Here, we used the Solanaceae family as a model to investigate differences in gene family size and the likely factors contributing to these differences. We found that genes in highly variable families have high turnover rate and tend to be involved in processes that have diverged between Solanaceae species, whereas genes in low-variability families tend to have housekeeping roles. In addition, genes in high-and low-variability gene families tend to be duplicated by tandem and whole genome duplication, respectively. This finding together with the observation that genes duplicated by different mechanisms experience different selection pressures suggests that duplication mechanism impacts gene family turnover. We explored using pseudogene number as a proxy for gene loss but discovered that a substantial number of pseudogenes are actually products of pseudogene duplication, contrary to the expectation that most plant pseudogenes are remnants of once-functional duplicates. Our findings reveal complex relationships between variation in gene family size, gene functions, duplication mechanism, and evolutionary rate. The patterns of lineage-specific gene family expansion within the Solanaceae provide the foundation for a better understanding of the genetic basis underlying phenotypic diversity in this economically important family.

## Introduction

Biological diversity can be attributed to the influence of the environment as well as genetic differences. One prominent source of genetic variation within and between species is gene copy number. Due to differential gene gains and losses, there can be substantial variation in the number of gene copies in a gene family, with some families exhibiting high turnover rates and others remaining similar sizes across species. In some cases genes in families with high turnover rates are involved in divergent biological processes (Tatusov, 1997; Rubin, 2000; Hahn et al., 2007; Guo, 2013). Thus, this high degree of turnover in gene family membership is expected to contribute significantly to divergence in cellular and developmental processes across species. Consequently, differences in gene family content can be shaped by natural selection (Pal et al., 2006; Schrider and Hahn, 2010) and are central to the evolutionary diversification and ecological adaptation of species (Demuth and Hahn, 2009; Zmienko et al., 2014; Carretero-Paulet et al., 2015). Thus, comparative studies of the patterns of gene family turnover are fundamental for understanding and assessing the functional, evolutionary, and ecological significance of duplicate genes.

In eukaryotes, gene duplication is the primary source of new genes that serve as the raw material for the evolution of novel functions (Ohno, 1970; Zhang, 2003). Duplicate genes can be generated through several mechanisms, such as whole genome duplication (WGD), segmental duplication, tandem duplication, and transposon-induced duplication (Panchy et al., 2016), each of which can have different impacts on duplicate gene functions and evolutionary fates and genomic architecture. WGD, for example, simultaneously doubles the number of all genes, and the requirement to maintain dosage balance leads to the preferential retention of genes encoding components of macromolecular complexes (Edger and Pires, 2009; Birchler and Veitia, 2014; Tasdighian et al., 2017). Segmental duplications in metazoans, where genomic segments that are hundreds to millions of base pairs long are duplicated in unlinked locations, can lead to chromosomal instability (Samonte and Eichler, 2002). Tandem duplication can lead to new gene structures through the formation of chimeric genes (Rogers et al., 2017) and contributes to preferential retention of genes involved in stress response (Hanada et al., 2008). Transposon-induced duplication generates duplicates such as retrogenes that are mostly dead on arrival (Brosius, 1991) but may modify gene expression (Flagel and Wendel, 2009).

Although duplicates can be preserved through acquisition of novel function(s) (neo-functionalization; Zhang, 2003), partitioning of ancestral functions among duplicates (sub-functionalization; Force et al. 1999), and/or other mechanisms (Lehti-Shiu et al., 2017), the majority of duplicate genes experience a brief period of relaxed selection and become pseudogenes within a few million years (Lynch and Conery, 2000). Because of differential gains, mostly due to differences in rates of gene duplication and loss through pseudogenization, gene family sizes and duplicate gene turnover rates are highly variable across species, including yeast (Hahn et al., 2005), fruit flies (Hahn et al., 2007), mammals (Demuth et al., 2006), and plants (Guo, 2013). The existing studies of gene family turnover in plants have focused on highly divergent taxa, ranging from green algae to flowering plants (Guo, 2013) or across the core eudicots (Carretero-Paulet et al., 2015). Thus, the extent of gene family size variation, and the factors, particularly gene duplication mechanisms and pseudogenization, that contribute to this variation among closely related plant species remain unclear.

Here we used the Solanaceae family as a model to investigate gene family variation among closely related species because a number of economically important species/cultivars in this family have been sequenced recently, including tomato (Tomato Genome Consortium, 2012), potato (Potato Genome Sequencing et al., 2011), eggplant (Hirakawa et al., 2014), pepper (Kim et al., 2014; Qin et al., 2014), tobacco (Sierro et al., 2013), and petunia (Bombarely et al., 2016). Fruits, tubers, leaves and flowers of these species/cultivars have been used by humans as food, medicine, stimulants and decoration. In addition, Solanaceae species are important models for functional characterization of plant genes (Vanden Bossche et al., 2013; Fan et al., 2016) and for evolutionary and ecological studies (Hu and Saedler, 2007; Nakazato et al., 2010; Särkinen et al., 2013). Furthermore, the genome sizes vary widely across Solanaceae species, ranging from 900 Mb in tomato (Tomato Genome Consortium, 2012) to 4.5 Gb in tobacco (Sierro et al., 2013). Through a comparative genomics analysis of 12 Solanaceae and 3 outgroup species, we first determined the number of domain family gains and losses in each lineage. Next we assessed the extent of variation in domain family size across species and the domain family turnover rate for each branch in the Solanaceae species phylogeny. Finally, we determined how gene duplication mechanisms and pseudogenization contribute to domain family size variability.

## Results & Discussion

### Domain family presence/absence variation

Variation in gene family size among taxa, which contributes to evolutionary divergence, is due to differential gain and loss of duplicates. Prior to assessing variation in gene family size, we evaluated the extent to which gene families were shared among species by examining the presence/absence distribution of gene families (using Pfam domains as a proxy, see Methods) across 12 Solanaceae species. In total, 4,313 families had ≥1 member in ≥1 species and were analyzed further. Of these domain families, 87.6% (3,775) were present in ≥10 species (**Fig. 1A**), suggesting they were present in the Solanaceae common ancestor, 2.9% of domain families (126) were present in 2-6 species, and 4.0% of domain families (174) were species-specific. To rule out the possibility that these lineage-/species-specific domain families are false negatives, we investigated three technical sources of error including: (1) missing annotations, (2) contamination during sequencing, and (3) genome assembly coverage and quality.

**Fig. 1.**
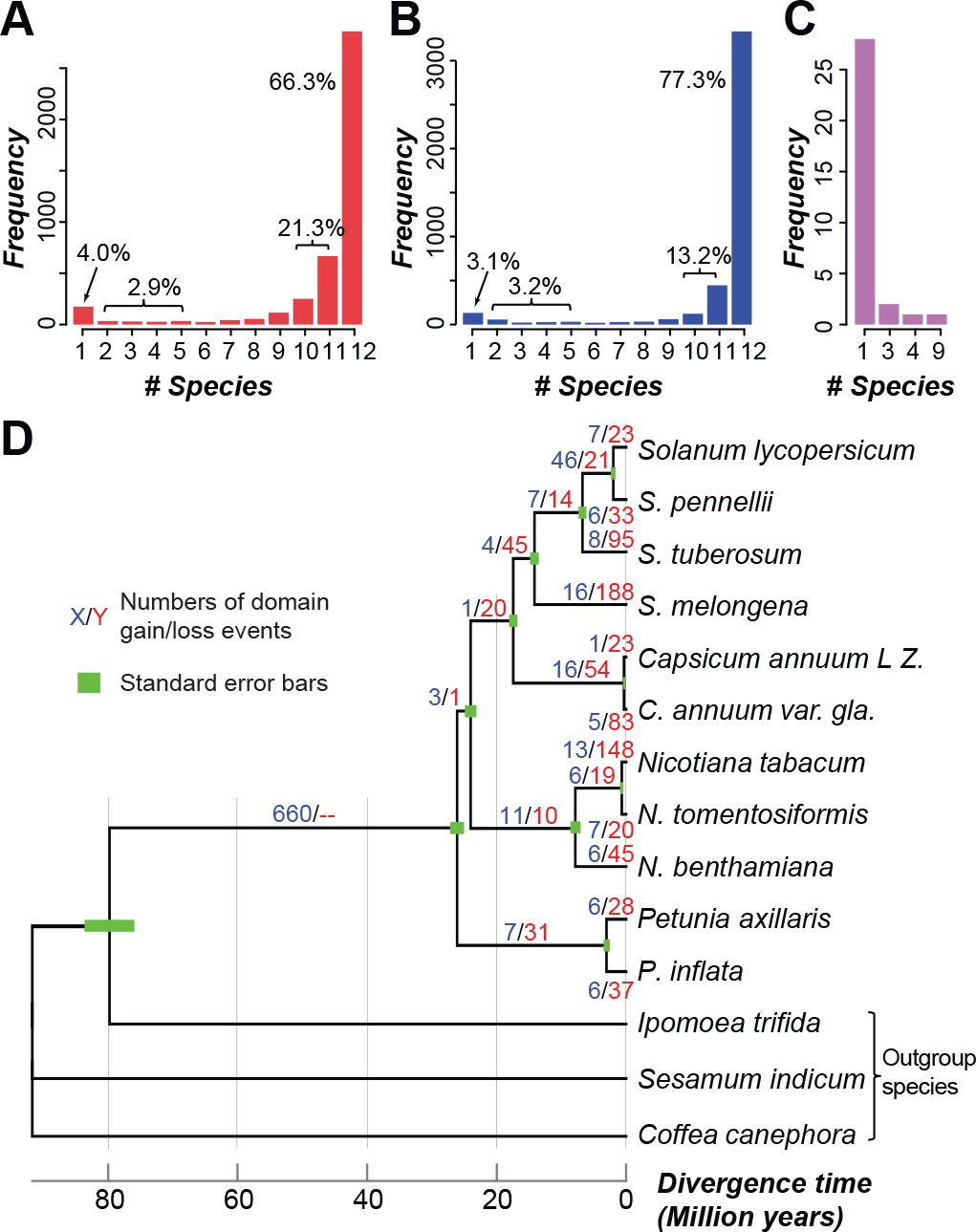
Distribution of domain families and domain family gains/losses in Solanaceae species. Frequency of domain families present in (A) the annotated genomic regions and (B) the annotated plus intergenic regions of different numbers of Solanaceae species. (C) The number of *N. sylvestris*-specific domain families (defined based on searches of annotated regions) present in intergenic regions of different numbers of Solanaceae species. (D) Inferred numbers of domain family gain/loss events across Solanaceae lineages. Blue/red numbers: the number of domain family gains and losses, respectively. Green bars: standard errors for divergence time estimates.

To determine if a domain family in a species was absent because it was not annotated, we used the seed sequences of each domain family to search against the intergenic sequences of that species. Out of 1,281 domain families that were absent in ≥1 species, sequences for 702 (54.8%) could be found in the intergenic regions of ≥1 other species (**Fig. 1B**). This indicates that annotation significantly impacts the number of domain families identified in a species. On the other hand, consistent with the contamination hypothesis, 72.4% of the species-specific domain families (126 of 174) were present only in *Nicotiana sylvestris*. Of the 126 *N. sylvestris*-specific domains, only 32 could be found in the intergenic regions of other species (**Fig. 1C**), further supporting the notion that some of these domain families may have been encoded by contaminating DNA introduced during sample collection or DNA extraction. Therefore, we excluded all *N. sylvestris* domain families and 41 other species-specific domain families that were not identified in the intergenic regions of any other species, leaving a total of 4,146 domain families.

Considering that the genomes we analyzed are of draft quality, we next determined the correlation between the scaffold N50 and the number of domain families absent in each species. We found no significant correlation (Pearson correlation coefficient [PCC]=-0.23, *p*=0.49), suggesting that even though incomplete genome assembly is expected to impact domain family discovery, the effect of this impact is not large enough to detect. Taken together, most domain families are present in nearly all Solanaceae species, indicating common ancestry. The existence of a subset of lineage/species-specific domain families, is largely explained by missing annotations, and 167 of these families appear to be derived from contamination.

### Inference of ancestral domain presence/absence states

With potential false negative cases identified and potential contaminating sequences removed, we next assessed the contribution of differential gains and losses to the lineage-specific distribution of domain families by inferring the ancestral presence/absence states of domain families in the 11 Solanaceae species (**Fig. 1D**). Of 757 domain families absent in ≥1 Solanaceae species, 660 and 71 were inferred to have been present and absent in the Solanaceae common ancestor, respectively, and 26 had ambiguous ancestral states (**Table S1-3**). To further assess the ancestral states of domain families in Solanaceae, we also analyzed the absence/presence distribution of domain families in other land plant/algae species (**Fig. 2**, **Table S3**). We found that 75% (73 of 97) of the domain families inferred to be absent or ambiguous based on analysis of Solanaceae species are present in multiple (>3) other plants/algae, indicating that these 73 families may also have been present in the Solanaceae common ancestor but had a higher loss rate compared with other domains. In addition to differential loss, the lineage-specific distribution of these families could also be due to high evolutionary rates where homologous domains are no longer recognized as belonging to the same domain family. To assess this possibility, we compared the *Ka*/*Ks* ratios of reciprocal best match gene pairs from *S. lycopersicum* and *S. pennellii* for domain families present in 2-11 species, regardless of ancestral state inference, and found no significant difference (Wilcoxon rank sum test, **Fig. S1**). This suggests that the lineage-specific distribution of domain families may not be significantly influenced by high evolutionary rates. Therefore, among the lineage-specific domain families inferred to exist in the common ancestor of Solanaceae species, most have likely been lost independently in ≥1 species.

**Fig. 2.**
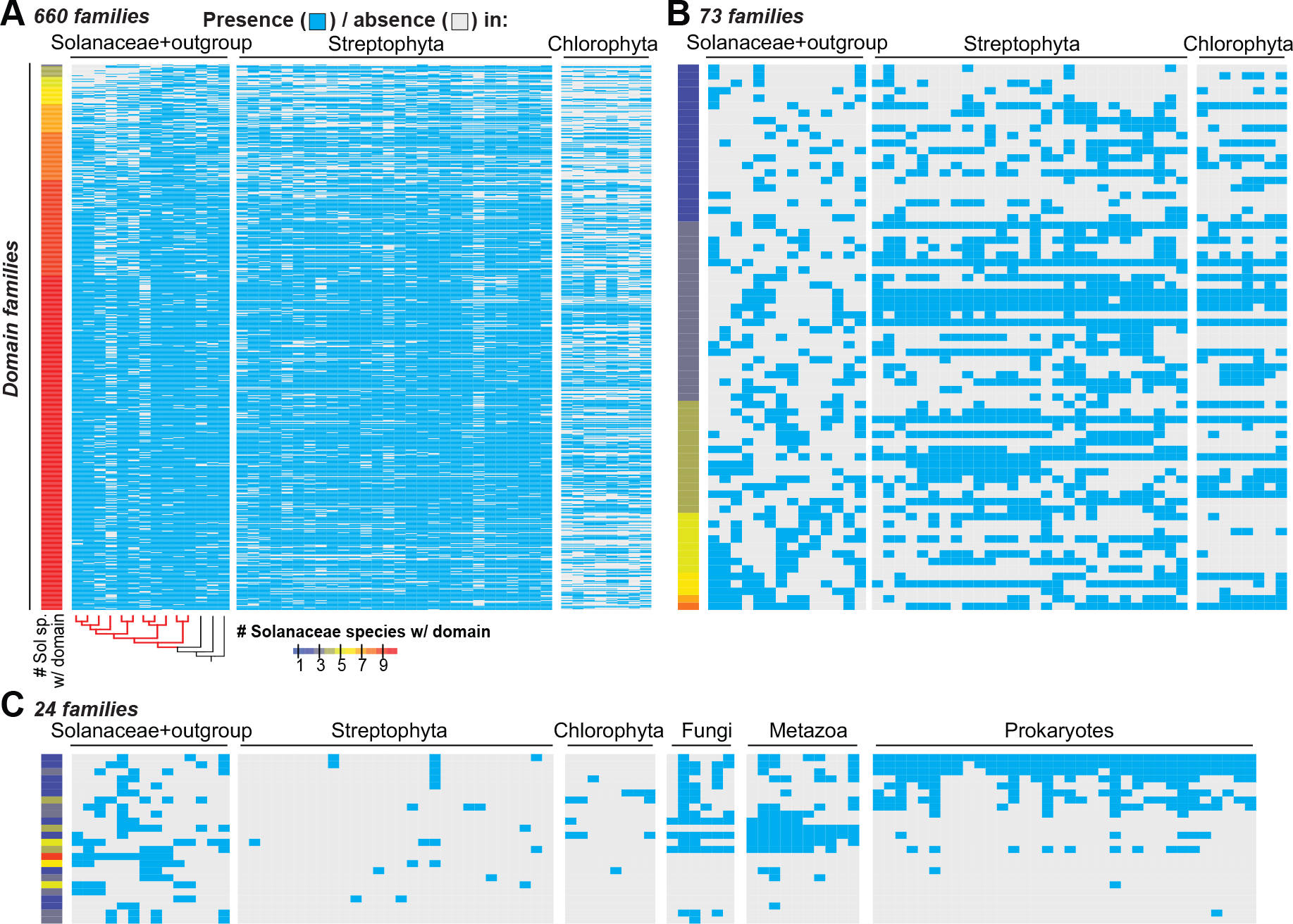
Distribution of domain families that are absent in ≥1 Solanaceae species among 11 Solanaceae species, 36 other plant/algal species, and 50 representative prokaryotic and eukaryotic phyla. Heat maps showing the presence/absence in each species for (A) 660 domain families inferred to be present in the Solanaceae common ancestor, (B) 73 domain families inferred to be absent in the Solanaceae common ancestor or that have an ambiguous presence/absence ancestral state but are present in >3 other plant/algal species, (C) 24 remaining domain families that are present in ≤3 other plant/algal species. Cyan: present; grey: absent. Color scale: the number of Solanaceae species with a given domain family. The phylogenic tree shows the same phylogenic relationships as Fig. 1D. Red and black branches indicate Solanaceae and outgroup species, respectively. Streptophyta and Chlorophyta species names and fungal, metazoan and prokaryotic phyla are shown in Table S3 and S4, respectively.

The remaining 24 families absent in ≥1 Solanaceae species and in most of the examined algal/plant species may have been: (1) present in the Solanaceae common ancestor but not identified in the Pfam hmmscan analysis based on the parameters we used (see Methods), (2) acquired due to *de novo* emergence of novel domains, (3) acquired through horizontal gene transfer (HGT), (4) contamination. Based on our Pfam hmmscan analysis in other plants and algal species, case (1) cannot be completely ruled out but is unlikely. We extended our analysis to examine the distributions of these 24 lineage-specific domains in 50 other representative prokaryotic and eukaryotic phyla (**Fig. 2C**). We found two families that are only present in a monophyletic group of closely related Solanaceae species but not in any other organisms examined. These include the Sar8_2 family, which is present only in non-*Petunia* species and has members involved in response to microbial infection (Alexander et al., 1992; Verberne et al., 2000), and the Prosystemin family, which is present only in *S. lycopersicum*, *S. pennellii* and *S. tuberosum*, and has members involved in wound response (Constabel et al., 1998). Although this may suggest the *de novo* origin of these two families, prosystemin represents a clear case of rapid divergence as structural homologs are also present in *Nicotiana* (Ryan and Pearce, 2003). We also found seven families that are present in monophyletic groups of Solanaceae species, but are also found in non-plant organisms (Fig. 2C **and Table S4**). Some of these domain families may have arisen through HGT, and this requires further analysis. In summary, the independent losses and gene annotation issues noted in the previous section are the two primary contributors to the limited distribution of some domain families, while other factors, such as *de novo* gains, rapid divergence, HGT, and contamination, likely only account for the limited distribution of a very small number of families.

### Variation in domain family size among Solanaceae species

After examining the distribution of domain families among Solanaceae species, we next assessed how the sizes of these families vary across species by measuring the coefficient of variation (CoV, standard deviation in domain family size divided by the mean size) of each domain family (**Table S5**). Because the mean domain family size is the denominator in CoV, similar degrees of changes in size have a greater effect for smaller families. To minimize this impact, we first binned the domain families based on their average sizes across species, determined the 95^th^ and 5^th^ percentile values of the CoV distribution for each bin, and fitted 95^th^ and 5^th^ percentile values across bins (**Fig. 3A**). Domain families above the 95^th^ and below the 5^th^ percentile trend lines were defined as having significantly higher and lower size variability compared with the genome-wide average, respectively. In total, there were 228 high-variability families and 410 low-variability families (**Table S5**).

**Fig. 3.**
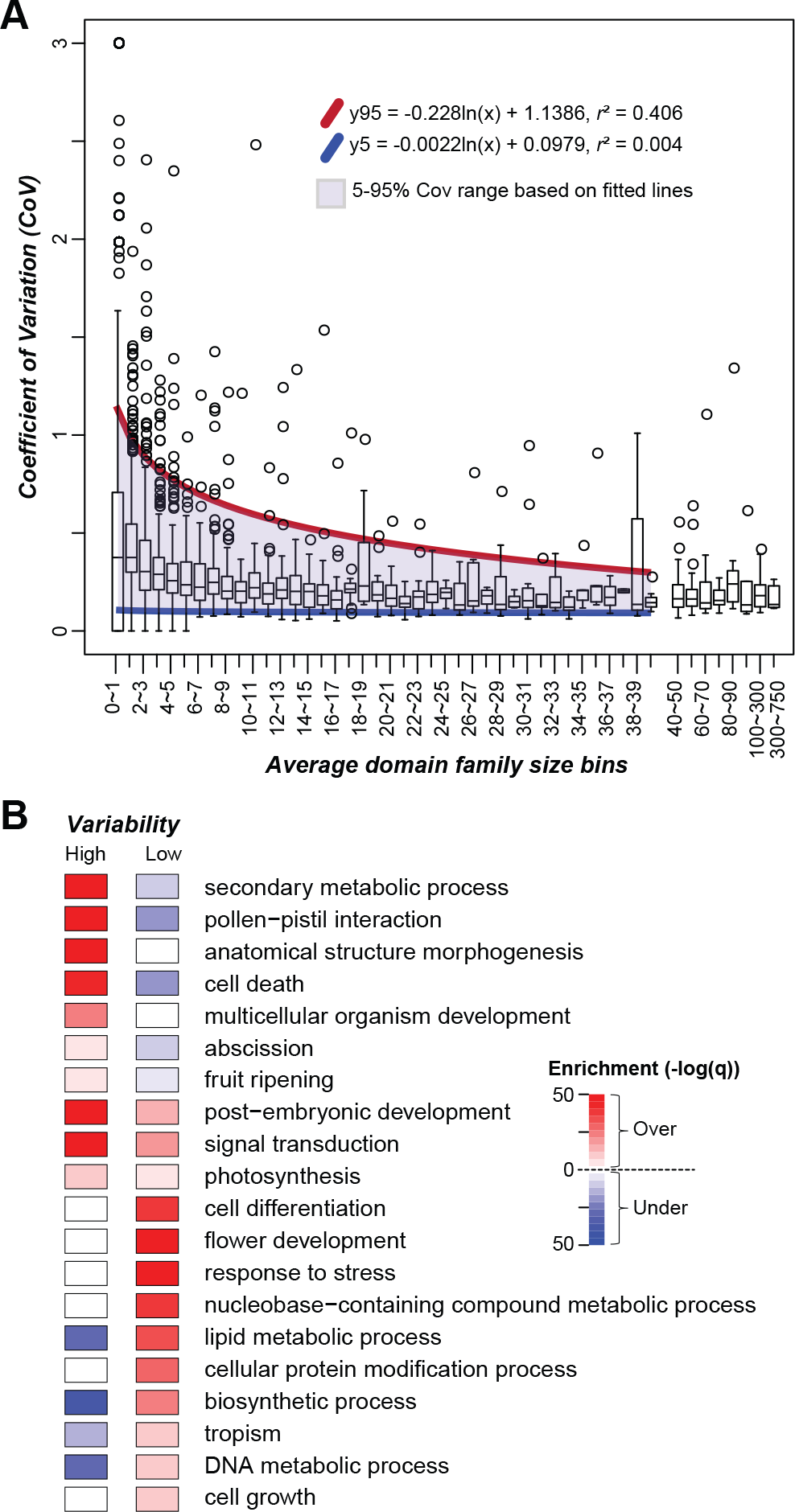
Relationship between domain family size (number of genes) and size variability.
(A) Distribution of the coefficient of variation (CoV) of domain family size among average domain family size bins. Domain families were assigned to bins based on the average number of genes in a domain family across species. Purple shade: region between the 5th (blue) and 95th (red) percentile trend lines. (B) GO Slim enrichment of genes in high- and low-variability domain families. Color scale: −log_10_(q-value). Red and blue: over- and under-representation, respectively.

To assess the functions that genes in high/low-variability domain families tend to have, we conducted a gene set enrichment analysis (see **Methods**). We found that genes in high-variability families tend to have functions related to, for example, secondary metabolic process, pollen-pistil interaction, cell death, abscission, and fruit ripening (**Fig. 3B**, **Table S6**,). An example of a high-variability domain family is 2OG-Fe(II) oxygenase (CoV=0.26, 84.6^th^ percentile), and diversification of 2OG-FeII-Oxy domain-containing genes is a key factor contributing to the diversity and complexity of specialized metabolites in land plants (Farrow and Facchini, 2014; Kawai et al., 2014). Another example is the NB-ARC domain (CoV=0.42, 100^th^ percentile), which is enriched in genes involved in cell death (De Oliveira et al., 2016). The eukaryotic protein serine/threonine/tyrosine kinase domain family (Pkinase, CoV=0.15, 66.7^th^ percentile) is also highly variable, and detailed analysis has shown that variability is mostly due to receptor-like kinases involved in self/non-self recognition (Lehti-Shiu and Shiu, 2012). The high CoV values observed for these families likely reflect rapid changes in response to the environment, particularly biotic factors.

In contrast, genes in domain families with low variability tend to be involved in central metabolism processes and housekeeping functions, including cell differentiation and growth, and lipid, protein and DNA metabolism (**Fig. 3B**). This indicates that negative selection contributes to low gene family variability. Surprisingly, genes in low-variability domain families tend to have functions in the response to stress, suggesting that some stress response processes may be consistently maintained across Solanaceae species. Genes from 4 to 38 domain families were annotated to each of the above GO categories, revealing how genes from different domain families interact to influence the underlying processes. For example, domain families enriched in genes related to tropism include PHY (Phytochrome) and two PHY-associated domains (HisKA and HATPase_c), as well as AUX_IAA. Phytochromes function as photoreceptors, while Aux/IAA genes regulate auxin-induced gene expression and also mediate light responses (Reed, 2001). Our observations are consistent with previous studies, showing the connection between light sensing and auxin signaling (Colon-Carmona et al., 2000; Halliday et al., 2009; Pedmale et al., 2010).

Taken together, domain family sizes vary considerably across species. The families with high size variability tend to be those that function in plant-environment interactions, particularly biotic interactions, where the high variability is likely a consequence of an evolutionary arms-race. In contrast, low-variability families tend to have housekeeping roles where strong negative selection has likely contributed to the maintenance of consistent family sizes across species.

### Gene gain and loss patterns among domain families

Domain family sizes can vary across species due to differences in domain family expansion or contraction rates in different lineages. To estimate how gene gain and loss events have contributed to the size variation of each domain family, we used a likelihood-based method (BadiRate, see **Methods**) to estimate the numbers of gene gain and loss events for internal and external branches in the Solanaceae species tree (**Fig. 4A**). The estimated average gene turnover (gain or loss) rate (λ) is 3.5e-2 events per gene per million years (MY), which is ~25 fold higher than an earlier estimate of λ across Viridiplantae (1.4e-3, including species from green alga to core eudicots spanning ~725 MY of evolution) (Guo, 2013). Because the Solanaceae species we included in our analysis span only ~26 MY of evolution, one possibility is that the shorter divergence time scale allowed us to better detect fluctuations in λ values that were masked across longer time scales (Demuth and Hahn, 2009). However, the Solanaceae λ is also ~17-30 fold higher than the turnover rates among yeast species (λ = 2.0e-3, ~32 MY) (Hahn et al., 2005), *Drosophila* species (λ = 1.2e-3, ~60 MY) (Hahn et al., 2007), and mammals (λ = 1.6e-3, ~93 MY) (Demuth et al., 2006).

**Fig. 4.**
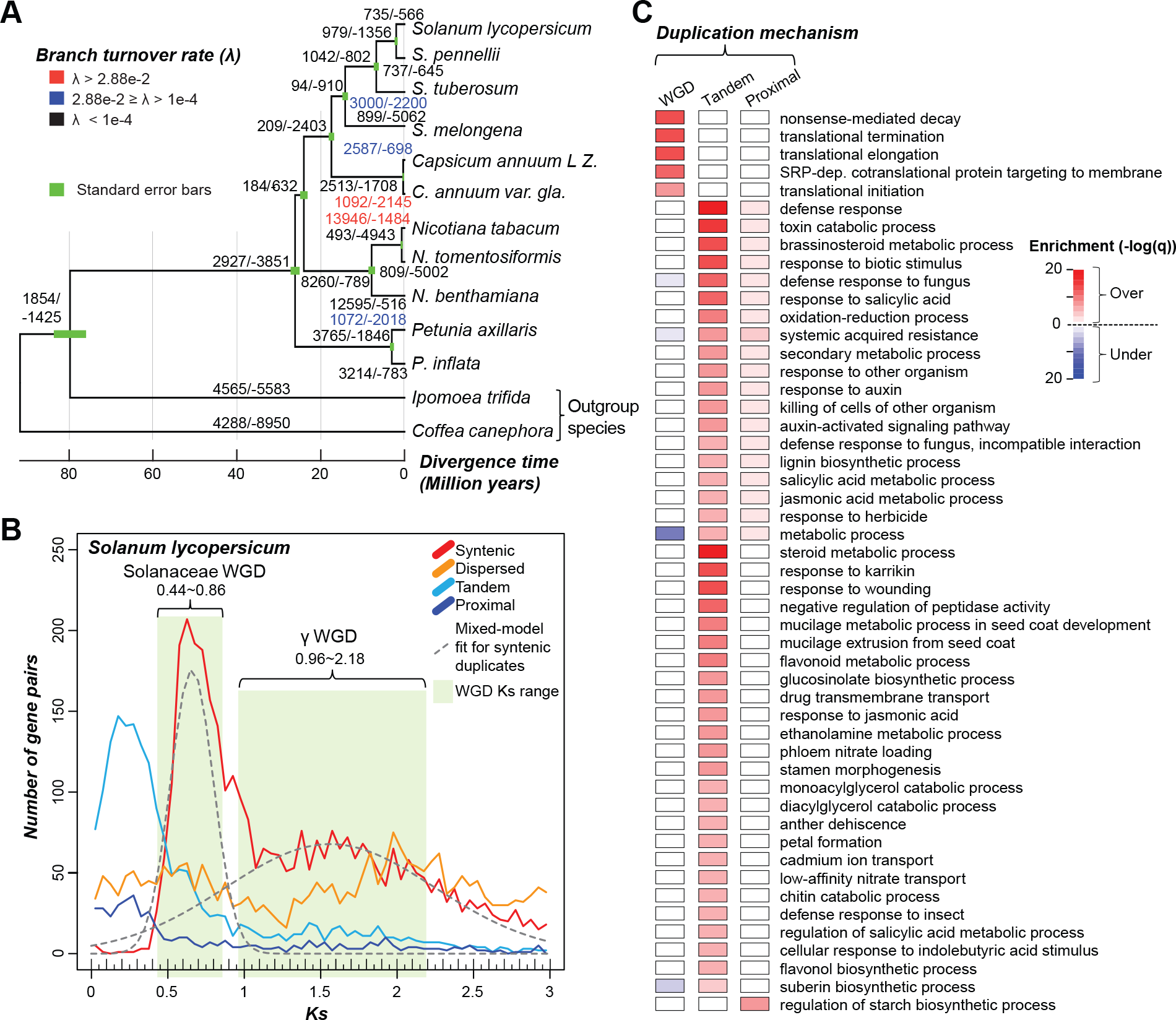
Gene gain and loss events and duplication mechanisms. (A) The total number of inferred gene gain/loss events across all families for each internal and external branch are shown. The numbers are colored based on inferred turnover rate (λ) as shown in the legend at the top-left corner. The phylogeny is the same as in Fig. 1D with *S. indicum* removed. (B) *Ks* distribution of *S. lycopersicum* duplicates derived from four different duplication mechanisms. Due to the high proportion of recent tandem duplicate genes and possible saturation of *Ks*, only duplicate genes with *Ks* values between 0.005 and are shown. Gray dashed lines show the fitted distribution of Sol and γ WGD duplicates. Means/ standard deviations (μ/σ) of the Sol and γ WGD distributions were 0.66/0.14 and 1.56/0.68, respectively. The ranges of *Ks* for the Sol and γ WGDs are indicated by green shading, and the cutoff values of *Ks* (μ±1.5σ and μ±0.9σ, respectively) are shown. (C) Enrichment of GO terms for *S. lycopersicum* genes duplicated by WGD, tandem and proximal duplications. Color scale: −log_10_(q-value). Red and blue: over- and under-representation, respectively.

Because divergence of these groups of species occurred on a similar time-scale as the Solanaceae species, another possibility is that the high Solanaceae λ is the consequence of recent large-scale duplication events. To assess this possibility, we more closely examined two branches with λ values larger than the average value. The first is the branch leading to *N. tabacum* (λ = 6.2e-1), which is derived from a recent allopolyploidy event, the hybridization of *N. tomentosiformis* and *N. sylvestris* (Sierro et al., 2013), and *N. tabacum* and *N. tomentosiformis* only diverged ~0.7 MYA (**Fig. 4A**). The second largest λ (4.0e-2) is on the branch leading to *C. annuum* var. *glabriusculum*, which was reported to have rapid amplification of transposable elements (Park et al., 2012; Qin et al., 2014). It is possible that the elevated transposable element activity may have led to more transposable element-mediated gene duplication events (Feschotte and Pritham, 2007; Freeling, 2009), resulting in a higher positive turnover rate. If these two highest λ values are removed, the average λ is 1.5e-3, similar to the gene turnover rate in Viridiplantae and other eukaryotes. Therefore, recent WGD and, to a lesser extent, transposon-mediated duplication, likely contributed to the significantly higher gene turnover rate among Solanaceae species.

The average λ varied not only between different branches, but also between different domain families. We hypothesized that high-variability domain families would have higher turnover rates. Consistent with this hypothesis, when the two branches leading to *N. tabacum* and *C. annuum* var. *glabriusculum* were excluded, the average λ for each domain family was significantly and positively correlated with the CoV percentile values (rho=0.43, *p*-value<2.2e-16). The average λ for high-variability domain families (1.9e-3) is multiple orders of magnitude higher than that for low-variability domain families (3.4e-8). This is also true if the *N. tabacum* and *C. annum* species are included (λ = 1.5e-2 and 1.3e-5 for high- and low-variability families, respectively). These findings indicate that, as expected, higher variability is the result of higher gene turnover.

### Influence of duplication mechanism on gene gains

Gene duplication and pseudogenization are two major factors leading to gene gains and losses, respectively. Because genes duplicated by different mechanisms are retained at different rates, we next assessed the extent to which different duplication mechanisms contributed to gene gains among Solanaceae domain families. To evaluate whether gene duplication mechanisms impact domain family size variation and gene turnover rate, we classified duplicate genes into four categories: (1) syntenic - duplicates in collinear blocks within the genome, which are likely derived from WGD or segmental duplication, (2) dispersed - duplicates located in unlinked locations but not in collinear blocks, (3) tandem - duplicates immediately adjacent one another, and (4) proximal - duplicates in close proximity but with intervening non-homologous gene(s) (see **Methods**). To determine when these duplication events took place, the synonymous substitution rate (*Ks*) between duplicates was used as a proxy for duplicate divergence time. In tomato for example, most tandem and proximal duplicate pairs have smaller *Ks* values than syntenic and dispersed duplicates, indicating they had a relatively more recent origin (**Fig. 4B**). This is also true for the other Solanaceae species (**Fig. S2**). The syntenic and dispersed duplicates were further divided into two bins, each corresponding to one of two rounds of WGD (Solanaceae-specific [Sol] and γ, see **Methods**).

Because gene retention is also influenced by gene functions (Zhang, 2003; Hanada et al., 2008; Edger and Pires, 2009), we next asked whether genes duplicated through different mechanisms tend to have different functions. For this analysis, we used *S. lycopersicum* as a representative species because it is the most extensively annotated among our target species. Genes duplicated by WGD tend to be involved in transcription and translation processes (Fig. 4C **and Table S7**), consistent with findings from earlier studies (Papp et al., 2003; Wu et al., 2008). In contrast, genes duplicated by tandem/proximal duplication tend to have functions in stress responses and secondary metabolic processes, which is also consistent with analyses of tandem duplicates in other plant species (Rizzon et al., 2006; Hanada et al., 2008).

The relative proportions of duplicates derived from different mechanisms and the patterns of *Ks* distribution vary greatly across species (**Fig. S2**). In particular, some species have either very few (e.g. *S. melongena*) or no syntenic duplicates (e.g. *N. tomentosiformis*). We found that assembly quality significantly influenced the discovery of syntenic duplicate genes, as genomes with smaller N50s tended to have fewer syntenic duplicates (rho=0.817, *p*=0.002). With this caveat in mind, we found that genes in 89.3%, 7.6%, and 3.1% of domain families were predominantly duplicated by WGD (includes syntenic and dispersed pairs that have *Ks* values corresponding to the Sol and γ WGDs; **Fig. 4B**), tandem, and proximal mechanisms (**Fig. 5A**, **Fig. S3**, **Table S8**). At one extreme, genes containing the Ribosomal_L5e domain were only duplicated by WGDs, while genes in the Sar8_2 family were only duplicated by tandem duplication. We next used Fisher’s exact test to determine whether members of a family were significantly more likely to be duplicated via a particular mechanism (**Table S9**). For DNA-binding transcription factor (TF) domain families (red text, **Fig. 5B**), we found that 11 TF families tended to be duplicated by WGD (all *p* < 5.1e-06), while only 3 TF families (SRF-TF, B3 and AP2) tended to be duplicated by tandem/proximal duplications (all *p* < 1.2e-06). Additionally, 18 predominantly primary metabolic enzyme domain families tended to be duplicated by WGD (all *p* < 5.2e-06), while 31 mostly specialized metabolic enzyme families (e.g. UDPGT and 2OG-FeII_Oxy) tended to be duplicated by tandem/proximal duplications (all *p* < 8.8e-06; **Fig. 5B**, **Table S9**). These results are consistent with studies postulating that TFs and primary metabolism gene duplicates are likely retained due to dosage balance requirements, whereas secondary metabolism genes have likely expanded lineage-specifically (Rizzon et al., 2006; Birchler and Veitia, 2007; Hanada et al., 2008; Freeling, 2009; Chae et al., 2014).

**Fig. 5.**
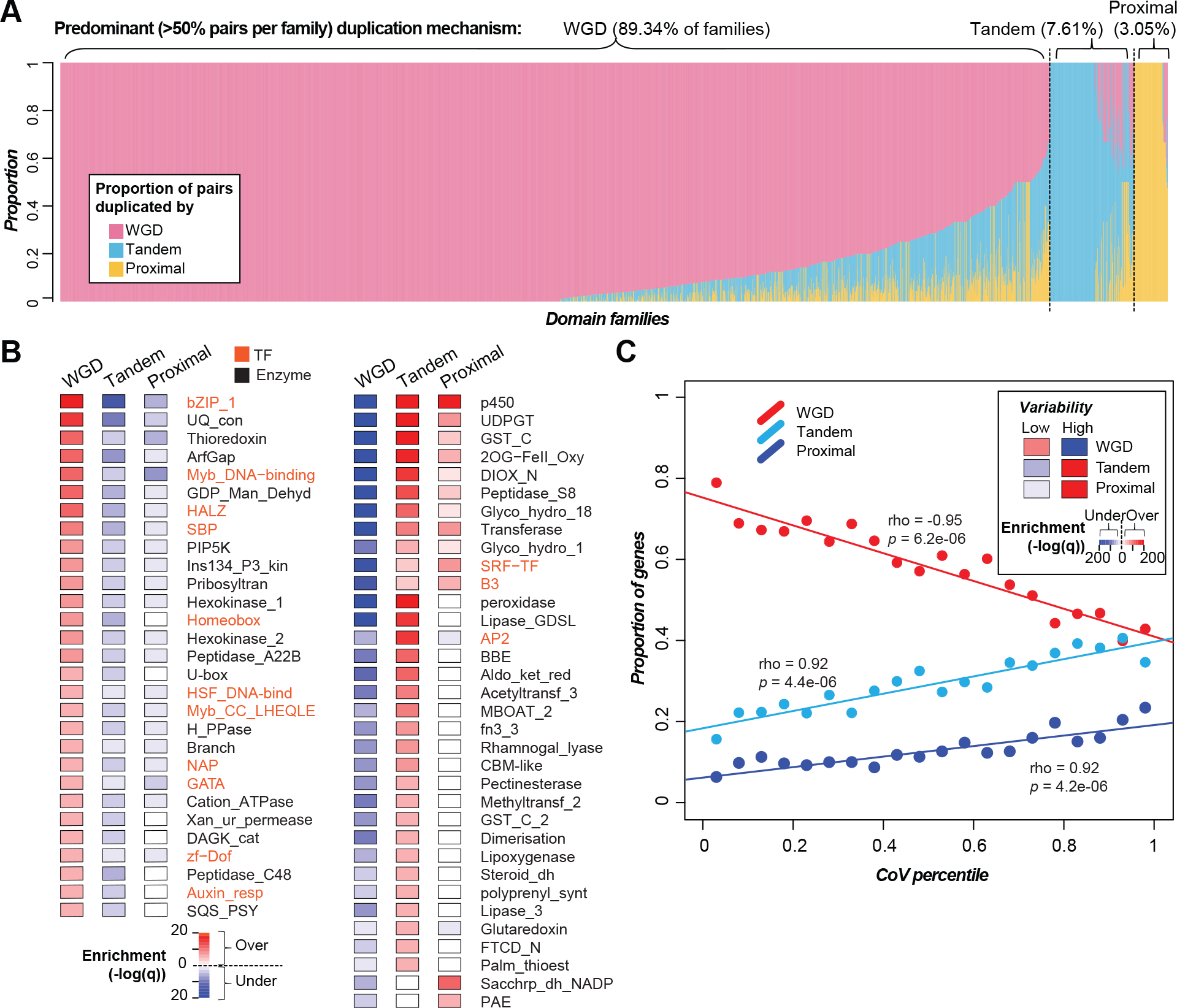
Contribution of duplication mechanism to domain family size variation in 11 Solanaceae species.
 (A) Proportion of duplicate pairs in each domain family (x-axis) that were predominantly duplicated by WGD, tandem, or proximal mechanisms. (B) Enrichment of members of DNA-binding transcription factor (orange) and enzyme (black) domain families that were duplicated via different mechanisms (full list in Table S7). Left panel: domain families that tend to be enriched in WGD duplicates. Right panel: families that tend to be enriched in tandem/proximal duplicates. (C) Correlation between domain family size variability (represented by CoV percentile) and the proportion of genes duplicated by different duplication mechanisms. The insert shows the enrichment of genes duplicated via differential mechanisms in high- and low-variability domain families tested with Fisher’s exact test. Color scale: −log_10_(q-value). Red and blue: over- and under-representation, respectively.

We next assessed the contribution of different duplication mechanisms to variation in domain family size. Because tandem/proximal duplications are more likely to be lineage-specific compared with WGDs, we expected and found that genes in high-variability domain families tended to be duplicated by tandem/proximal duplications (cyan and blue lines, **Fig. 5C**). In contrast, the proportion of WGD duplicates is anti-correlated with CoV percentile (red line, **Fig. 5C**). Consistent with these observations, genes in high-variability domain families (above the 95^th^ percentile trend line, **Fig. 3A**) tended to be duplicated by tandem and proximal duplication, while genes in low-variability domain families (below the 5^th^ percentile trend line) tended to be duplicated by WGDs (insert, **Fig. 5C**). These patterns highlight the fact that different duplication mechanisms contribute differently to gene gains among Solanaceae domain families in a way that exhibits substantial functional bias. In particular, tandem/proximal duplications are the main contributor to lineage-specific differences. In contrast, the finding that WGD duplicates tend to be enriched in low-variability domain families suggests that these duplicates are consistently either retained or lost post-duplication among different lineages.

### Relationship between the timing and mechanism of duplication and selective pressure

As described in previous sections, variability in gene family size is strongly correlated with duplication mechanism and gene function. We next asked if Solanaceae genes duplicated by different mechanisms have significantly different evolutionary rates based on the ratio of non-synonymous substitution rate (*Ka*) to *Ks* of each duplicate pair. To capture duplication events more thoroughly, the duplicates examined included both annotated genes and pseudogenes in the 11 Solanaceae species. Thus, we focused on three types of duplicate pairs: (1) GG: gene-gene, (2) GP: gene-pseudogene, and (3) PP: pseudogene-pseudogene pairs that were further classified based on the potential duplication mechanisms (**Fig. S4 and Fig. 6**).

**Fig. 6.**
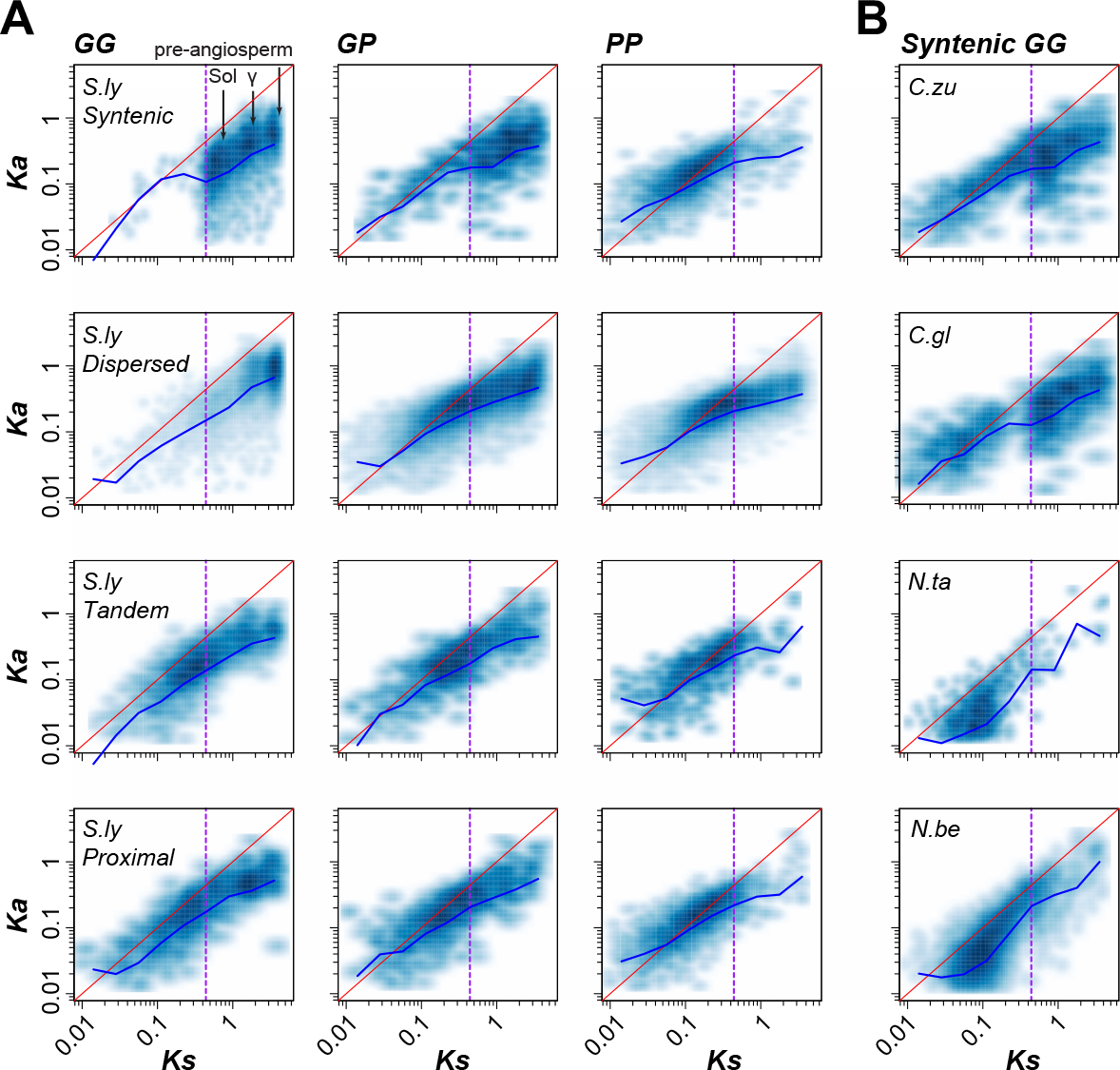
Evolutionary rates of different duplicates in representative species. (A) Gene-gene (GG), gene-pseudogene (GP) and pseudogene-pseudogene (PP) pairs duplicated by different mechanisms (syntenic, dispersed, tandem and proximal) in *S. lycopersicum* (*S.ly*). Three high density regions in the *S.ly* syntenic GG *Ka*-*Ks* plot correspond to the Sol, γ and pre-angiosperm WGDs, respectively. (B) Syntenic GG pairs in *C. annuum* L. *zunla*-1 (*C.zu*), *C. annuum_var. glabriusculum* (*C.gl*), *N.tabacum* (*N.ta*) and *N. benthamiana* (*N.be*). Each point in a *Ka*-*Ks* dot plot represents a single pair of duplicate sequences, and darker blue denotes a higher density of points. Red lines indicate the expectation under neutral selection, and blue lines connect the median *Ka* value of each log_10_ (*Ks*) bin as shown in Fig. S5B and C. The vertical purple dashed line shows the lower boundary (*Ks* = 0.44) for defining duplicates derived from the Sol WGD.

In *S. lycopersicum* for example, there were significantly fewer syntenic (43.7%) and tandem (43.3%) GP/PP duplicate pairs than dispersed (81.6%) and proximal (84%) duplicates (Fisher’s exact tests comparing the numbers of GG and GP/PP pairs, all *p*-values < 2.2e-16), which may indicate that dispersed and proximal duplicates are more likely to become pseudogenes, and thus, may be evolving faster. Consistent with this interpretation, syntenic GG pairs had the lowest *Ka/Ks* values, followed by dispersed, tandem and proximal GG pairs (Wilcoxon signed-rank test, all *p*-values < 6.1e-5, Fig. 6A **and Fig. S5A**). One potential reason why tandem GG pairs had higher *Ka/Ks* values than dispersed GG pairs is that most tandem GG pairs were duplicated more recently (**Fig. 4B and Fig. 6A**), and younger duplicates tend to experience more relaxed selection (Lynch and Conery, 2000). For each duplication mechanism, the *Ka/Ks* values tended to be the highest for GG pairs, followed by GP and PP pairs. This is expected given that pseudogenes should evolve neutrally. What is surprising is the presence of PP pairs with high *Ks* but low *Ka/Ks* ratios (the third column, **Fig. 6A**). The low *Ka/Ks* ratios indicate that the pseudogenization of these PP pairs likely occurred relatively recently because the signature of past selection remains. Thus, these PP pairs are examples of duplicate pairs that persisted for a long period of time (tens of millions of years) but eventually became pseudogenes.

As expected, a high proportion of syntenic GG pairs are likely derived from WGD (referred to as WGD pairs), and three high density regions in a plot of GG *Ka* vs. *Ks* values correspond to the Sol, γ and pre-angiosperm WGDs, respectively (**Fig. 6A**). Only 1.4% of syntenic GG pairs had *Ks* < 0.44 (lower bound for defining the Sol WGD) and are likely derived from recent segmental duplication (referred to as segmental pairs). We also found that the *Ka/Ks* values of WGD GG pairs (0.17 on average) were significantly lower than those of more recently duplicated, segmental GG pairs (0.47 on average) (Wilcoxon signed-rank test, *p*-values = 3.76e-13, Fig. 6A **and Fig. S5B**). These observations indicate that recent segmental GG pairs may evolve faster than WGD GG pairs and tend to become pseudogenes quickly. To rule out the impact of divergence time on evolutionary rate of duplicate genes (Lynch and Conery, 2000), we compared two *C. annuum* cultivars with a large number of recent, segmental GG pairs (*Ks* < 0.44) against two *Nicotiana* species (*N. tabacum* and *N. benthamiana*) with a recent WGD (**Fig. 6B and Fig. S5C**). We found that *C. annuum* segmental duplicates had significantly higher *Ka/Ks* values than *Nicotiana* WGD duplicates (Wilcoxon signed-rank test, all *p*-values < 2.2e-16). Therefore, recent segmental GG pairs have higher *Ka/Ks* even when divergence time is taken into account. Taken together, these results suggest that genes duplicated by different mechanisms experience different selection pressures, consistent with a previous study (Yang and Gaut, 2011), which eventually results in different rates of gene retention and loss.

### Contribution of pseudogenization to variation in domain family size

On average, 20.9% and 52.0% of duplicate pairs in *S. lycopersicum* are GP and PP pairs, respectively (**Fig. S4**). This indicates that gene loss occurred frequently. Therefore, we expect that gene loss significantly contributes to variability in domain family sizes among Solanaceae species. Although variability in domain family size is significantly correlated with pseudogene number, the correlation is weak (Spearman’s rank correlation coefficient, rho=0.15, *p*<2.2e-16, **Fig. 7A**). We also found that species with more genes do not necessarily have more pseudogenes (rho=0.1, *p*=0.78, **Fig. 7B**). For example, although *N. tabacum* and *N. benthamiana* experienced the most recent WGD and have the largest number of protein-coding genes, they don’t have the largest number of pseudogenes (**Fig. 7B**). Instead, there is a significant positive correlation between pseudogene number and genome size (rho=0.81, *p*=0.003, **Fig. 7C**), consistent with the hypothesis that a larger genome size is likely the consequence of less efficient removal and/or more frequent expansion of ‘non-functional’ sequences (Lefebure et al., 2017). We also found that larger domain families tend to have more pseudogenes (**Fig. 7D**), indicating that these domain families tend to experience both more frequent gene birth and death events.

**Fig. 7.**
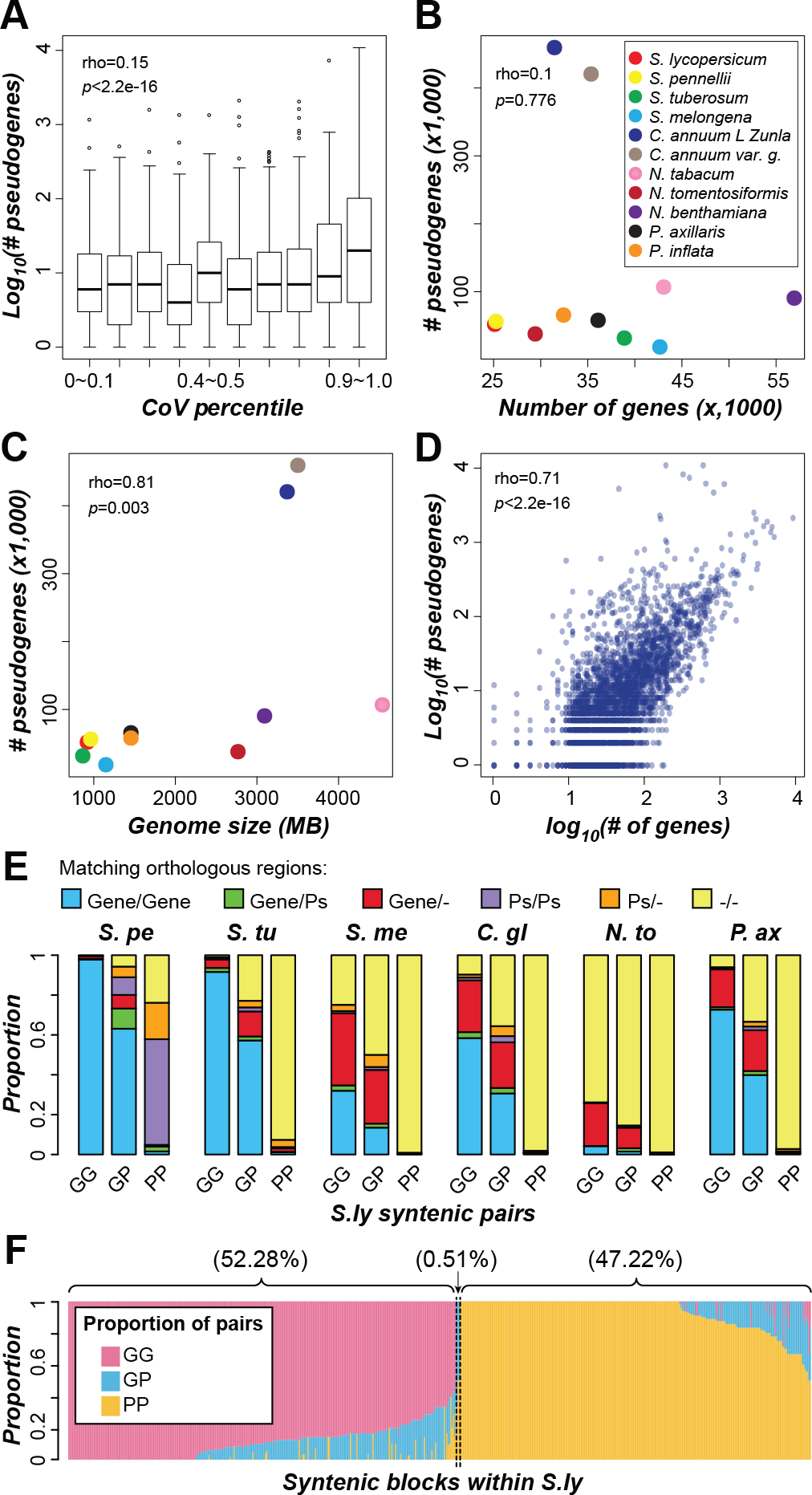
Contribution of pseudogenization to domain family size variation. (A) Relationship between domain family size variability (CoV percentile) and logarithmic number of pseudogenes. (B) Relationship between the number of genes and pseudogenes among Solanaceae species. (C) Correlation between genome size and number of pseudogenes. (D) Correlation between the logarithmic number of genes and pseudogenes in a domain family. Each dot indicates a domain family. The rho and p value for Spearman’s rank correlation are shown. (E) Proportion of different *S. lycopersicum* (*S.l*y) syntenic duplicate pairs that have orthologous sequences in collinear regions in other species. The orthologous sequences were defined giving priority to protein-coding genes over pseudogenes. If orthologous protein-coding genes were not identified for a given gene, then orthologous pseudogenes were searched for. *S.pe*: *S. pennellii*, *S.tu*: *S. tuberosum*, *S.me*: *S. melongena*, *C.gl*: *C. annuum_var. glabriusculum*, *N.to*: *N. tomentosiformis*, *P.ax*: *Petunia axillaris*, Ps: pseudogene, “-”: no orthologous sequence was found. (F) Proportion of duplicate pairs in *S. lycopersicum* collinear blocks that are predominantly (>50%) GG, GP or PP pairs. Each column represents a pair of collinear blocks within *S. lycopersicum.* GG: gene-gene pair, GP: gene-pseudogene pair, PP: pseudogene-pseudogene pair.

In the previous section, we discussed PP pairs that are likely derived from WGD events but become pseudogenes independently (PP column, **Fig. 6A**). We also noted the presence of a substantial number of PP pairs derived from more recent duplication events (*Ks*<0.44, **Fig. 6A**). These recent PP pairs may be derived from independent pseudogenization of originally functional duplicates or may be duplicates of pseudogenes. To distinguish between these possibilities, we searched for orthologs of *S. lycopersicum* pseudogenes in six other Solanaceae species. We found that, out of 2,011 recent segmental PP pairs with *Ks* < 0.44, 1,885 (93.7%) (**Fig. 7E**) have either pseudogenes as orthologs or no apparent orthologous sequences in the syntenic regions of all Solanaceae species analyzed. This proportion (93.7%) is significantly higher than that observed for GG (0.4%) and GP (13.3%) pairs (Fisher’s exact test, both *p*<2.2e-16) and is inconsistent with the expectation that, if both sequences in a PP pair were pseudogenized independently after duplication, the corresponding functional homologs should be found in ≥1 other Solanaceae species. Thus, most recent segmental PP pairs are likely derived from pseudogene duplication, rather than pseudogenization after duplication of functional genes.

The origin of recent segmental PP pairs by duplication is also supported by the high proportion of pseudogenes in collinear, duplicated blocks within species. Among 593 pairs of collinear blocks in *S. lycopersicum*, 310 (52.28%) and 280 (47.22%) have predominantly GG and PP pairs, respectively (**Fig. 7F**); which is significantly deviated from the random expectation (z-scores 7.4 and 63.4, respectively; *p* values < 6.8e-14). Thus, collinear blocks tend to contain either GG or PP pairs. This pattern supports the notion that pseudogenes in these blocks are products of pseudogene duplication because, to explain this pattern based on independent pseudogenization of functional ancestral genes, a large number of additional, independent loss events would be required. The observation that recent segmental PP pairs tend to be located in regions with a high density of repeats (**Fig. S6**) suggests that these duplicates may be derived from repeat-mediated duplication mechanisms.

## Conclusion

Genomes of an increasing number of plant species have been sequenced, facilitating comparative studies aimed at evaluating genome and gene content evolution among related plant species and, specifically in this study, identifying factors contributing to gene family size variation. Here we show that the distribution of domain families across Solanaceae species varies due to lineage-specific gains or losses and that different duplication mechanisms have contributed to gene family size variation. Genes in domain families with higher variability are more likely to have been duplicated by tandem duplication. Most of the observed tandem duplicates were duplicated recently and tend to be involved in processes that are highly diverse among Solanaceae species, e.g., secondary metabolism (Chowański et al., 2016) and fruit ripening (Knapp, 2002). Genes duplicated though different mechanisms also have different evolutionary rates. For example, tandem and recent segmental duplicate genes experience more relaxed selection than WGD duplicate genes. Taken together, these findings suggest that lineage-specific gene family expansion through tandem duplication plays an important role in the evolution of organisms and diversification among closely related species. Comparative evolutionary and functional analysis (e.g., of gene structures and expression patterns) of new tandem or segmental duplicate genes and ancestral genes, may help to uncover genetic changes underlying lineage-specific innovations.

The abundance of pseudogenes is often used to estimate the extent to which gene loss has impacted gene family size (Demuth and Hahn, 2009). However, we found that pseudogenes are also frequently duplicated and remain readily detectable just like functional genes, indicating that the number of pseudogenes is not an accurate proxy for gene loss. Pseudogene duplication may happen randomly, producing duplicates that are not under selection. Alternatively, some pseudogene duplicates may be retained due to their effects on, for example, regulation of their protein-coding relatives (Pink et al., 2011). Further studies will be necessary to distinguish between these possibilities. We found that recent segmental PP pairs are closely associated with repeat sequences. It remains to be determined whether these recent segmental PP duplications in Solanaceae were produced by a recombination-like transposable element-mediated mechanism, as in humans (Zhou and Mishra, 2005), or by another yet to be discovered mechanism.

## Materials and Methods

### Genome annotation and domain family designation

The genome sequences and annotations for each Solanaceae species and three outgroup species were downloaded from National Center for Biotechnology Information (NCBI, https://www.ncbi.nlm.nih.gov/genome/) or Solanaceae Genomics Network (solgenomics, https://solgenomics.net/): *S. lycopersicum* V2.5 (NCBI), *S. pennellii* SPENNV200 (NCBI), *S. tuberosum* V3.4 (solgenomics), *S. melongena* r2.5.1 (solgenomics), *Capsicum annuum* L. *zunla-1* V2.0 (solgenomics), *C. annuum_var. glabriusculum* V2.0 (solgenomics), *N. tabacum* TN90 NGS (solgenomics), *N. tomentosiformis* V01 (NCBI), *N. sylvestris* GCF_000393655.1 (NCBI), *N. benthamiana* V1.0.1 (solgenomics), *Petunia axillaris* V1.6.2 (solgenomics), *P. inflata* V1.0.1 (solgenomics), *Ipomoea trifida* V1.0 (NCBI), *Sesamum indicum* V1.0 (NCBI), and *Coffea canephora* Vx (solgenomics). For two species with no annotation GFF files (*S. melongena* and *I. trifida*), CDS sequences were obtained from NCBI and used as queries in searches against the respective genome sequences using BLAST-like alignment tool (BLAT) (Kent, 2002) to find the location of each coding region in the genome. The criteria used for mapping were a threshold of 100% identity and no gap tolerated if located within an aligned block. The GFF files were then generated based on the BLAT output with blat2gff.pf (Kent, 2002). To identify pseudogenes, protein sequences from *A. thaliana*, *O. sativa* and *S. lycopersicum* were used as queries in tblastn (Altschul et al., 1990) searches against the genome sequences of target species, and intergenic sequences with significant similarity to known proteins were further processed with the pipeline from Campbell et al. (2014). The chromosome locations of protein-coding genes and pseudogenes were drawn with Circos (Krzywinski et al., 2009).

Pfam domain Hidden Markov Models (HMMs, Version.3.0) were downloaded from the Pfam database (Finn et al., 2014), and transposase domains or domains found in proteins with transposase domains were excluded from downstream analyses. Protein sequences of genes in each Solanaceae species were used as queries in searches against the Pfam HMMs using HMMER3 (Finn et al., 2011) with the trusted cutoff. If >1 domains overlapped, the overlapping region was annotated with the Pfam domain with the smallest E-value. All protein sequences containing the same Pfam domain were considered to be in the same domain family. Genes with >1 type of protein domain were counted as being members of each domain family, thus a single gene can belong to multiple domain families. For comparison, the domain family sizes in eukaryotic species in representative phyla were downloaded from the Pfam database (Finn et al., 2014). To avoid the confounding effects of WGD in coefficient of variation calculations, domain family sizes from *N. tabacum* and *N. benthamiana*, which have experienced recent WGDs (Bombarely et al., 2012; Sierro et al., 2013), were excluded.

### Species tree and ancestral presence/absence state inference

To build the species tree, genes in domain families with only a single copy in each species, and one randomly chosen copy if there were >1 copies in *N. tabacum* or *N. benthamiana*, which have experienced recent WGD (Bombarely et al., 2012; Sierro et al., 2013), were used. For each domain family, amino acid sequences were aligned using MUSCLE (Edgar, 2004), and poorly aligned regions were removed using trimal (Capella-Gutiérrez et al., 2009) with a gt cutoff value of 0.8, (i.e. columns with gaps in more than 20% of the sequences are removed). The alignments were then combined and used to build a phylogenetic tree using RAxML/8.0.6 (Stamatakis, 2014) with the following parameters: -f a -x 12345 -p 12345 -# 1000 -m GTRGAMMA, with sequences from *C. canephora* set as outgroups.

To estimate the species divergence times, fourfold degenerate transversion rates (4DTv) were used for the Molecular Clock Test in MEGA (Tamura et al., 2013), using the General Time Reversible model and Gamma Distributed (G) of rates among sites. The evolutionary rate was set as 6×10^−9^ per site per year (Wolfe et al., 1989). The estimated species divergence times based on the molecular clock are consistent with an earlier study (Särkinen et al., 2013). The ancestral presence/absence states of domain families were first inferred using the parsimony method (**Table S1**) in Mesquite (Maddison and Maddison, 2017). For nodes with ambiguous states, the maximum likelihood method was used to choose the more likely states (**Table S2**).

### Functional annotation

Protein sequences were used as queries in blastp searches against the NCBI nr protein database with an E-value cut-off of 1e-5, and Gene Ontology (GO) annotations were inferred using blast2go (Conesa and Götz, 2008) with default parameters. Some genes may be annotated with GO terms related to non-plant activities, e.g., GO:0001568 (blood vessel development); therefore, GO terms from *Arabidopsis thaliana* (http://www.arabidopsis.org/), *Oryza sativa* V7.0 (http://genome.jgi.doe.gov/) and *S. lycopersicum* ITAG2.4 (ftp://ftp.solgenomics.net/) annotations were used as reference lists to filter out non-plant GO terms. Plant GO Slim terms (http://www.geneontology.org/) were used to obtain a broad overview of functional annotation. Gene set enrichment analysis was performed using Fisher’s exact test, and the *p*-value was adjusted to account for multiple testing (Benjamini and Hochberg, 1995). GO terms with adjusted *p*-values smaller than 1e-5 were considered significantly over/under-represented.

### Domain family turnover rate and sequence evolutionary rate calculation

We used the likelihood-based method implemented in BadiRate v1.35 (Librado et al., 2012) to estimate the rate of gene gains and losses (turnover rate λ). Three different branch models including Free Rates (FR, each branch has its own turnover rate), Global Rates (GR, all branches have the same turnover rate), and Branch-specific Rates (BR, particular branches have specific turnover rates), were used to estimate λ values. To take into account potential differences in gene turnover rates due to overall gene family expansion and rapid gene loss after lineage-specific WGDs (Sankoff et al., 2010; Inoue et al., 2015), in the BR model, all the branches leading to lineages with WGDs were assigned branch-specific turnover rates, while other branches were assumed to have the same turnover rate. For large domain families, where no results were obtained after a >100 hour runtime, Markov Clustering (Enright et al., 2002) was used to divide each gene family into smaller subfamilies with the parameter −I=3. The best turnover rate model for each domain family was chosen based on likelihood ratio tests (Peers, 1971).

To determine which Solanaceae lineages have experienced WGD events, all-against-all blastp searches were performed both within and between species to obtain within and between species reciprocal best match protein-coding gene pairs, respectively. Evolutionary rates were then determined by calculating the synonymous (*Ks*) and non-synonymous (*Ka*) substitution rates for gene pairs using the yn00 program in PAML (version 4.4.5) with default parameters (Yang, 2007). The *Ks* distribution of gene pairs within and between species was used to infer the relative timing of WGD events and speciation, respectively (**Fig. S7**). Consistent with earlier studies (Leitch et al., 2008; Potato Genome Sequencing et al., 2011; Tomato Genome Consortium, 2012; Hoshino et al., 2016), these *Ks* distributions indicated that the Solanaceae species experienced the γ triplication (γ WGD) shared by stem lineages of core eudicots (Vekemans et al., 2012), and the Solanaceae-specific triplication (Sol WGD). *N. tabacum* and *N. benthamiana* independently became polyploids after the Solanaceae-specific triplication (Bombarely et al., 2012; Sierro et al., 2013).

### Gene duplication mechanisms

To classify duplicate genes into different categories according to duplication mechanism, the software MCScanX-transposed (Wang et al., 2013) was used. First, based on intra-and inter-species all-against-all blastp results, the top five best-matching sequences to each query sequence with *p*-value ≤ 1e-10 were assumed to be homologs and retained. Then the chromosomal locations of these homologous genes were compared using MCScanX-transposed. For a given species, all other species in the same genus, one representative species from each of the other Solanaceae genera, and two non-Solanaceae species were used as reference species. For example, when gene duplication mechanisms were assessed for *S. lycopersicum* genes, gene sequences and chromosomal locations from *S. pennellii*, *S. tuberosum*, *S. melongena*, *C. annuum_var. glabriusculum*, *N. tomentosiformis*, *P. axillaris*, *I. trifida* and *C. canephora* were used in MCScanX-transposed.

Duplicate genes were classified into four categories: 1) syntenic duplicates - paralogs are located in corresponding collinear blocks within species; 2) dispersed duplicates - one paralog and its ortholog are both located in corresponding interspecies collinear blocks, while the other paralog is not; 3) tandem duplicates - paralogs are immediately adjacent to each other; 4) proximal duplicates - paralogs are adjacent each other but separated by ≤10 non-homologous genes. Duplicates that did not belong to any of the above categories were removed from our analysis as their mechanism of duplication is ambiguous.

Note that instead of using the category names in MCScanX, we renamed the “segmental duplicates” as “syntenic duplicates”, because genes in alignable blocks within each genome could have been duplicated by either WGD or segmental duplication (Cannon et al., 2004; Singh et al., 2015). We also renamed the “transposed duplicates” as “dispersed duplicates” because these genes could have been duplicated through transposition, WGD and subsequent rearrangement of one of the copies, recombination between repeat sequences in unlinked regions, or non-homologous end-joining of double-stranded breaks (Woodhouse et al., 2010). Because the *Ks* distributions of both syntenic and dispersed duplicates showed two similar peaks (**Fig. S2**), corresponding to the Sol and γ WGD events (**Fig. S7**), we further assigned them to these two WGDs, by estimating the *Ks* value distribution as a mixture of Gaussian distributions:

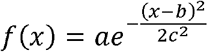

where a, b and c are fitted constants obtained using nonlinear (weighted) least-squares estimation (nls) of the *Ks* distribution in the R environment (Nash, 2014). After values of a, b and c were fitted for both WGD duplicate categories, distributions of simulated *Ks* were obtained. Cutoff values of *Ks* to define the boundaries of these two WGD events were chosen based on two criteria: 1) to maximize the difference in the area under curve between the two distributions (i.e. choosing cutoff values yielding the largest difference in area under the two fitted distributions); 2) for any given *Ks* value, the number of gene pairs from the distribution corresponding to the Sol WGD is > twice than the number with *Ks* values corresponding to the γ WGD distribution. Because the Sol and γ WGDs were experienced by all Solanaceae species and the synonymous substitution rate is often assumed to be neutral, we used the *Ks* cutoff values from *S. lycopersicum*, which has the genome assembly with the highest N50, to define WGD duplicate gene pairs in other species.

## Acknowledgments

We thank John P. Lloyd and Christina B. Azodi for helpful discussions. This work was supported by National Science Foundation IOS-1546617 and DEB-1655386 to S.-H.S.

